# The *Drosophila* IR20a Integrates L-arginine and Salt Signals via Distinct Subunit Assemblies

**DOI:** 10.64898/2026.05.28.728318

**Authors:** Bo Wang, Guangnan Qu, Xutong Wang, Hubert Amrein, Yan Chen

## Abstract

Amino acids and sodium salt are essential nutrients that frequently occur together in natural food sources. While animals typically employ distinct receptor subfamilies for specific taste modalities, the molecular mechanisms that integrate these concurrent nutritional signals at the peripheral level remain poorly understood. Here, we identify the *Drosophila* Ionotropic Receptor IR20a as a multimodal tuning receptor required for the detection of both the amino acid arginine and low NaCl salt. We find that IR20a is expressed in a subset of IR76b-positive gustatory receptor neurons (GRNs) distinct from GRNs dedicated to sweet or bitter taste. Mutation analysis and heterologous expression experiments demonstrate that IR20a functions combinatorially: it pairs with the core receptor IR25a to detect arginine, whereas the additional recruitment of IR76b is required to confer sensitivity to low salt. Crucially, we establish that IR20a/IR25a complex mediates synergistic responses to arginine-salt mixtures, thereby enhancing feeding preference. Furthermore, we reveal that IR20a provides a tonic, state-independent nutrient signal that operates in parallel to the starvation-modulated IR56b pathway to regulate NaCl feeding. Together, our studies uncover a molecular logic in which combinatorial assembly of receptor complexes enables the peripheral integration of complex nutritional signals.

## Introduction

The ability to sense and discriminate between chemical components in food is fundamental for the fitness and survival of all animals ^1^. Species across diverse phyla, including vertebrates and arthropods, exhibit conserved appetitive responses to essential nutrients like sugars, while generally demonstrating avoidance of potentially toxic bitter compounds ^1–3^. For amino acids and salt (NaCl), behavioral responses are often concentration-dependent: low concentrations of these key nutrients are attractive, whereas high concentrations frequently elicit aversion ^4–8^.

In mammals, the perception of the five basic taste modalities (sweet, sour, bitter, salty, umami) is largely thought to adhere to a ℌlabeled lineℍ coding strategy ^1,9,10^. In this model, specific taste modalities are mediated by distinct taste receptor cells expressing respective taste receptor - proteins^1,10^. In contrast, the gustatory system of the fruit fly, *Drosophila melanogaster*, is organized in a more complex fashion. While sweet and bitter tastes are generally encoded by distinct populations of gustatory receptor neurons (GRNs), individual GRNs can exhibit multimodal sensitivity ^11–13^. For instance, attractive nutrients such as sugars, low salt and some amino acids are sensed by sweet-sensing neurons, whereas chemicals to be avoided, such as bitter compounds, high salt, high acid, and certain amino acids are detected by bitter-sensing GRNs ^12–14^. Furthermore, water, acids, and aversive high concentrations of salt are detected by yet other GRNs, distinct from both sweet and bitter GRN populations ^15–17^. This multimodal nature is conferred by two types of chemoreceptors, often co-expressed within the same GRNs: about 60 Gustatory Receptors (GRs), which primarily encode receptors for sweet and bitter tasting chemicals, and about 64 Ionotropic Receptors (IRs), which are critical for sensing salts, acids, fatty acids and amino acids ^11–13,18^. GR as well as IR based taste receptors are multimeric complexes, most likely hetero tetramers composed of two to four different GRs and IRs, respectively. Most IR based complexes include IR25a and IR76b, which are largely co-expressed and thought to function as core subunits. They are complemented with one or two additional IR subunits that confer specificity for different types of ligands (referred to as tuning receptors) ^7,19–22^. For example, in sweet GRNs, IR25a and IR76b form complexes with IR56b to mediate low salt detection ^23–25^ and with IR56d to mediate fatty acid responses ^20,26^. In contrast, in bitter GRNs, IR25a and IR76b associate with IR51b to detect several amino acids ^7^ and with and IR94e to sense glutamate, respectively ^27^. And in a third set of GRNs expressing the pheromone receptor Ppk23 and the neurotransmitter glutamate (named Ppk23^glut^), IR25a and IR76b combine with IR7c to form a receptor for high concentrations of monovalent salts ^28^. These observations established that in addition to the multimodal nature of GRNs (expressing both GRs and IRs), specific tuning receptors combine with core subunits to form tetrameric IR complexes that are dedicated to the detection of subgroups of chemical ligands.

The natural environment presents food chemicals as mixtures rather than single-modality isolates. For instance, low salt and amino acids naturally co-occur in the fruit fly’s preferred food sources: rotting, yeast-fermented fruits ^29^. Consequently, taste systems must be capable of integrating signals from different taste modalities to guide foraging and feeding decisions ^13^. This integration occurs in the suboesophageal ganglion (SOG) and the Nucleus of the Solitary Tract (NTS), the primary taste processing center of insects and vertebrates, respectively, and plays a critical role in coordinating nutrient consumption ^13,30^. Furthermore, signal integration has been reported at the peripheral taste cell level ^31,32^. Specifically, while the labeled-line logic largely prevails in mammalian taste cells, significant ℌcross-talkℍ occurs within the taste bud ^32^. For example, ATP released from sweet- or bitter-sensing Type II cells can activate serotonin release in neighboring Type III cells, effectively processing and integrating information before it reaches afferent nerve fibers ^33,34^. Similarly, in flies, bitter compounds and high salt solutions suppress sweet responses at the level of GRNs within the same taste sensilla, whereas acids can alleviate bitter-induced suppression of sweet taste ^19,35,36^. However, despite these advances in understanding cellular and circuit-level processing, the possibility of integration occurring directly at the level of the gustatory receptor complex has remained largely unexplored.

In this study, we report that the IR20a functions as a multimodal tuning receptor essential for sensing both the amino acid arginine and low NaCl salt. We demonstrate that this dual sensitivity is achieved through the combinatorial assembly of a receptor complex: IR20a requires the co-receptor IR25a for arginine detection while the additional recruitment of IR76b is critically for low salt detection. Crucially, the IR20a/IR25a complex confers higher responses to amino acid-salt mixtures than each of the ligands alone. Heterologous expression of IR20a and IR25a in the pheromone sensing neurons and S2 cells reveals that this enhancement is synergistic rather than additive, indicating that the increased feeding preference arises from receptor synergy. Finally, we show that the IR20a pathway provides a distinct, state-independent layer of nutrient regulation that operates in parallel to the known starvation-modulated salt sensing pathway. Together, our findings uncover a fundamental molecular logic where the combinatorial architecture of peripheral receptor complexes provides a mechanism for integrating distinct, yet co-occurring, nutritional signals.

## Results

### *IR20a-GAL4* labels a novel population of GRNs responding to amino acids and salts

Previous studies have implicated IR20a in amino acid taste in tarsi and the modulation of salt sensitivity in the labellum ^37^. However, this study did not identify the specific tarsal GRNs for amino acid detection, nor did it establish whether IR20a is endogenously expressed in salt-sensing neurons to modulate salt taste in labellar GRNs. To clarify the cellular function of IR20a, we characterized its expression in the gustatory system and the brain using an *IR20a-GAL4* driver ^37^. Despite prior reports that IR20a is absent from the labellum ^38^, our confocal imaging uncovered approximately 18 weakly labeled GRNs in this organ (Fig. 1A), corroborating a proposed role for IR20a in high-pH detection ^39^. Beyond the labellum, IR20a marked two tarsal GRNs on each of the fore-, mid-, and hindlegs (Fig. 1B). In the pharynx, we detected two GRNs in the labral sense organ (LSO) and two in the ventral cibarial sense organ (VCSO) (Fig. 1D3 - D4), consistent with earlier findings ^38^; however, no signal was evident in the brain or ventral nerve cord (Fig. 1C). Co-expression analysis using *IR20a-GAL4*, IR25a antibodies and *IR76b-QF* revealed that IR20a is co-expressed with both IR25a and IR76b (Fig. 1D - E). However, we found no overlap between *IR20a-GAL4* and *GR64f-LexA* or *GR66a-LexA* (Fig. 1F - G). Thus, IR20a is co-expressed with the core receptors IR25a and IR76b in a population of neurons that is distinct from both sweet and bitter GRNs.

**Figure 1.**
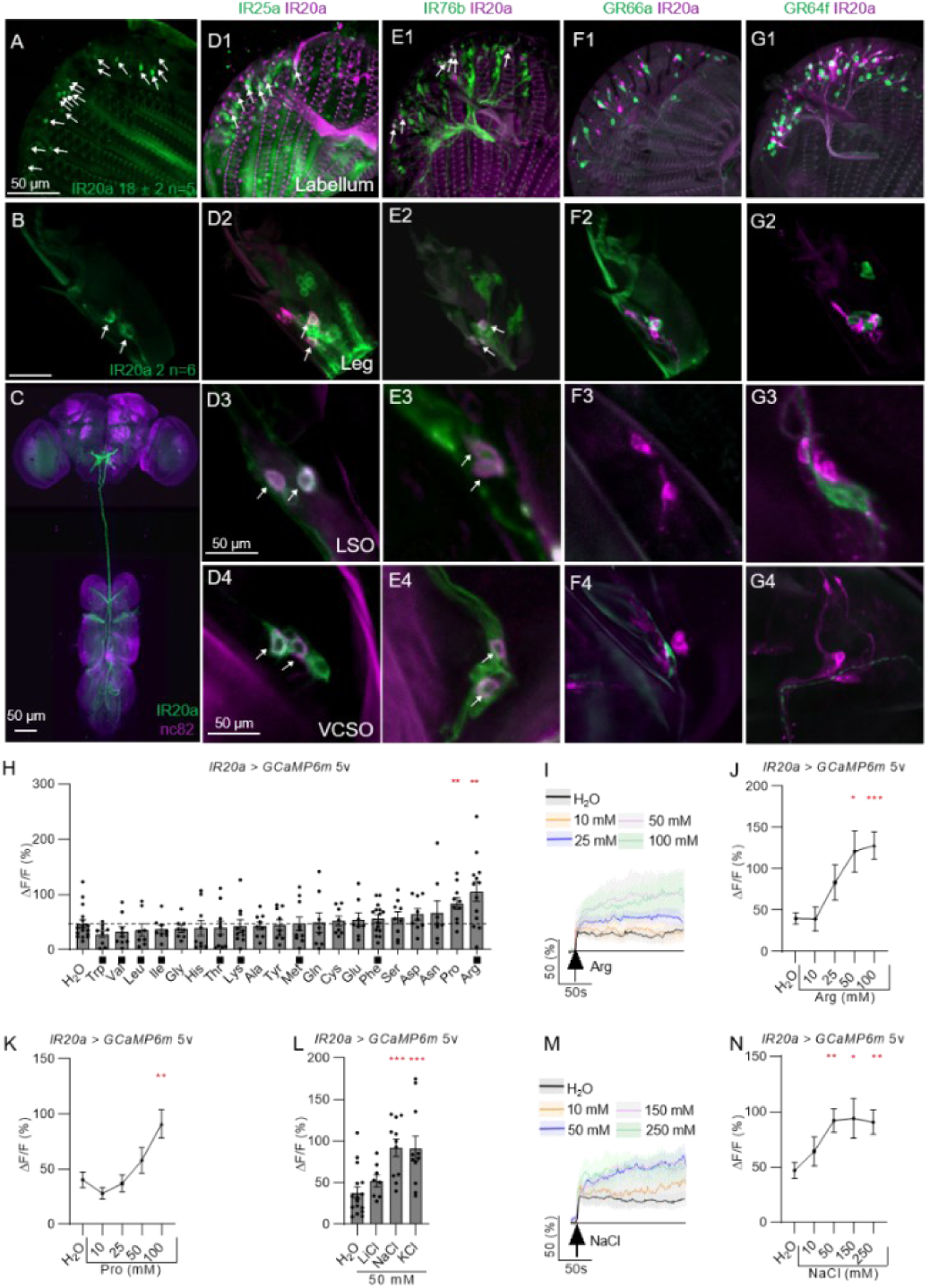
*IR20a-GAL4* labels a population of gustatory receptor neurons (GRNs) sensitive to specific amino acids and salts. **(A - B)** *IR20a-GAL4* expression is observed in approximately 20 GRNs in the labellum **(A)** and 2 GRNs in the fifth tarsal segment **(B)**; arrows indicate the cell bodies labeled by *IR20a-GAL4*. (C) *IR20a-GAL4* does not label cell bodies within the brain or ventral nerve cord (VNC); however, neuronal projections from the labellum, legs, and wings are visible in these central tissues. **(D - G)** Double-labeling experiments in the labellum, tarsi, and pharynx show that *IR20a-GAL4* is co-expressed with the co-receptors IR25a **(D)** and IR76b **(E)**. No overlap is observed with markers for bitter-sensing (*GR66a-LexA*) **(F)** or sweet-sensing (*GR64f-LexA*) **(G)** GRNs. Arrows indicate co-expression. Note that pseudocoloring was used for IR20a and IR76b in (E) for consistent visualization. **(H)** Single-cell calcium imaging of *IR20a-GAL4* tarsal GRNs reveals robust responses to 100 mM L-arginine and L-proline, but not to other basic amino acids. Note that black squares under the bars indicate essential amino acids for flies. **(I - K)** Dose-response analysis of *IR20a-GAL4* tarsal GRNs stimulated with 10 - 100 mM L-arginine **(I - J)** and 10 - 100 mM L-proline **(K)**. (L) *IR20a-GAL4* tarsal GRNs respond to 50 mM NaCl and KCl, but show no significant response to 50 mM LiCl. (M - N) *IR20a-GAL4* tarsal GRNs exhibit plateaued calcium responses to NaCl at concentrations of 50 mM and higher. Data represent mean ± SEM (n = 8 - 18). Mann-Whitney test comparing ligands to H_2_O control, *p < 0.05, **p < 0.01, ***p < 0.001.

To characterize the response profiles of IR20a-expressing GRNs, we performed single-cell calcium imaging of tarsal GRNs by expressing GCAMP6 under the control of *IR20-GAL4* ^40^. We focused on tarsal neurons because *IR20a-GAL4* labels GRNs very weakly in the labellum, precluding reliable detection of GCaMP signals. Based on previous reports implicating IR20a in amino acid detection ^37^, we first examined responses to 19 proteinogenic amino acids (Fig. 1H). Calcium imaging revealed that stimulation with 100 mM L-arginine or L-proline elicited robust calcium responses in IR20a GRNs, whereas other amino acids failed to induce significant activity (Fig. 1H). Dose-response analyses showed that only high concentrations of L-arginine (50 and 100 mM) and L-proline (100 mM) evoked statistically significant responses (Fig. 1I–K). We next examined salt sensitivity and found that IR20a tarsal GRNs responded strongly to 50 mM NaCl and KCl, but not to LiCl (Fig. 1L). NaCl-evoked responses were concentration dependent and reached a plateau at approximately 50 mM (Fig. 1M–N). We note that lysine was excluded from imaging experiments due to its intrinsic color, which interferes with fluorescence measurements; instead, its role was assessed behaviorally using the proboscis extension reflex (PER) assay in wild type and *IR20a* mutant flies, which revealed that IR20a is not required for lysine attraction (Fig. S1A - B).

Together, these results indicate that tarsal *IR20a* GRNs are multimodal sensory neurons that respond to both specific amino acids (arginine and proline) and monovalent salts (Na⁺ and K⁺).

### IR20a is required for arginine and low NaCl salt attraction

To determine whether IR20a is functionally required for cellular responses to amino acid arginine, proline, and monovalent salt Na^+^ and K^+^, we examined *IR20a* mutants using calcium imaging. Ca^2+^ responses to L-arginine, L-proline and KCl were preserved and not statistically different from those of wild type flies (Fig. 2A - C). In contrast, Ca^2+^ responses to 50 mM NaCl were completely abolished in *IR20a* mutants, a defect fully rescued by reintroducing *UAS-IR20a* (Fig. 2A - C).

**Figure 2.**
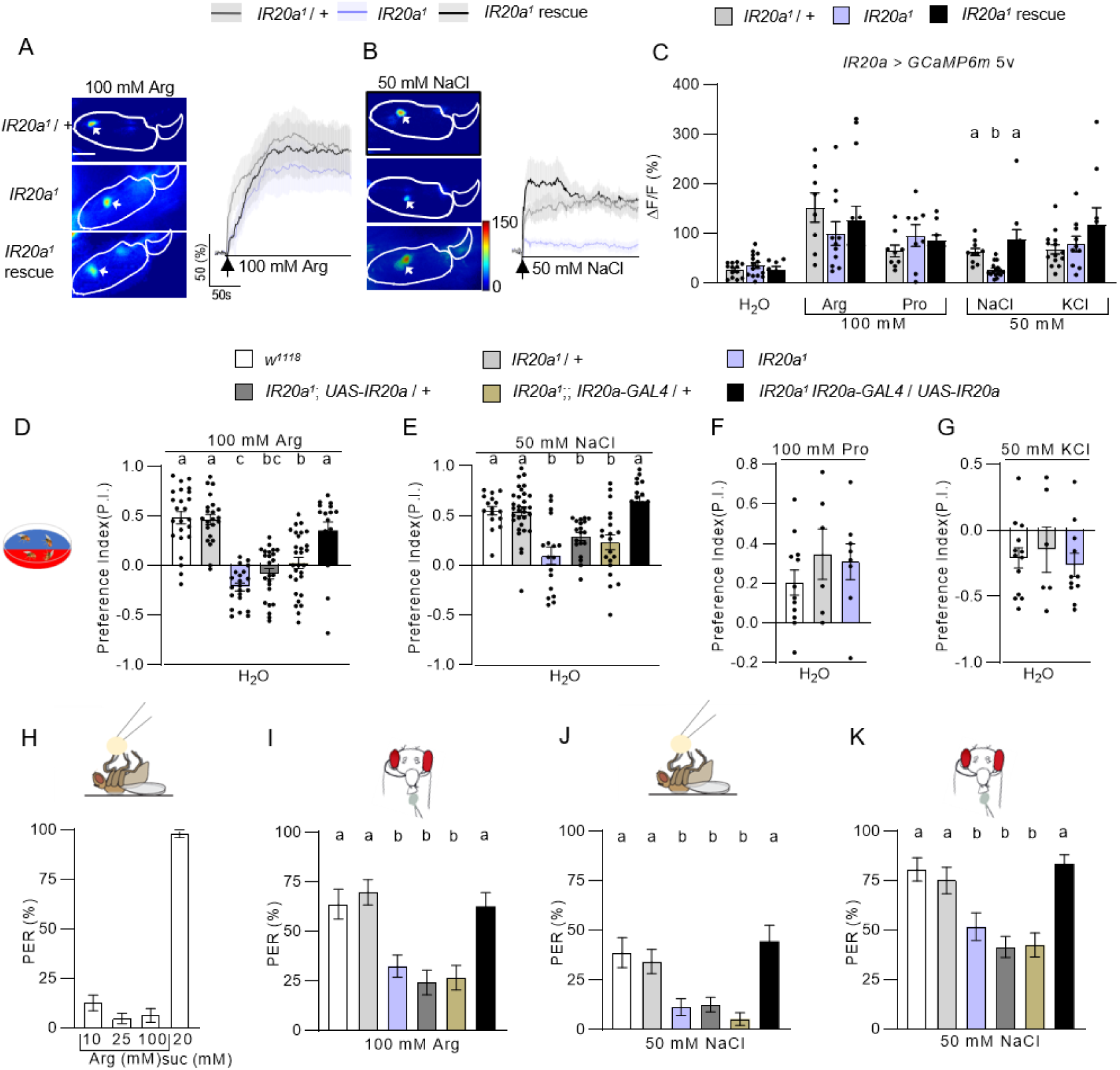
*IR20a* is required for low NaCl attraction and contributes to arginine-driven feeding behavior. **(A - C)** Single-cell calcium imaging reveals that the response of *IR20a-GAL4* tarsal GRNs to 50 mM NaCl is abolished in *IR20a* mutants, while responses to 100 mM L-arginine, L-proline, and 50 mM KCl remain intact. (n = 8 -16). **(D - E)** Behavioral attraction to 100 mM Arg **(D)** and 50 mM NaCl **(E)** is restored upon genetic rescue of *IR20a*. (n = 16 - 31). **(F - G)** IR20a is dispensable for preference toward 100 mM proline (Pro) **(F)** and 50 mM KCl **(G)**. (n = 18 - 39). **(H - I)** Tarsal stimulation with arginine does not induce PER **(H)**, but labellar stimulation with 100 mM arginine induces PER in an IR20a-dependent manner **(I)**. (n = 17 – 34). **(J - K)** *IR20a* is required for PER induced by 50 mM NaCl, with stimulation in the tarsal or labellum **(K)**. (n = 18 - 49). Data represent mean ± SEM. Mann-Whitney test **(H),** Kruskal-Wallis with Dunn’s post hoc test **(C - G and I - K).** Different letters denote differences between designated groups (p < 0.05)

We next assessed the behavioral relevance of IR20a using two-choice feeding assays. Although our imaging data indicated that IR20a is only necessary for detecting low concentration of NaCl but not arginine, *IR20a* mutants exhibited significantly reduced attraction to both 100 mM arginine and 50 mM NaCl, a phenotype that was fully rescued by including *UAS-IR20a* driven by *IR20a-GAL4* (Fig. 2D - E). In contrast, attraction to 100 mM proline and avoidance of 50 mM KCl were unaffected in *IR20a* mutants (Fig. 2F - G).

We further examined the role of IR20a using the proboscis extension reflex (PER) assay. Consistent with previous reports, tarsal stimulation with arginine failed to elicit PER in wild-type flies ^41^ (Fig. 2H). In contrast, labellar stimulation with arginine induced a strong PER that was markedly reduced in *IR20a* mutants and fully rescued by re-expression of IR20a (Fig. 2I). Together with the two-choice feeding assays, these results establish that IR20a is essential for arginine attraction. To determine whether IR20a is selectively required for arginine sensing, we examined PER responses to all other amino acids using distinct amino acid pools. No significant differences were observed between wild-type controls and *IR20a* mutants for any other amino acid, demonstrating that IR20a specifically mediates arginine attraction rather than amino acid attraction in general (Fig. S1C - D). We next tested PER responses to salt. Wild-type flies displayed weak PER upon tarsal stimulation and robust PER upon labellar stimulation with 50 mM NaCl (Fig. 2J - K). Both responses were significantly reduced in *IR20a* mutants and were restored by *IR20a-GAL4*–driven IR20a expression (Fig. 2J - K). Together, these findings indicate distinct roles for IR20a in arginine detection between labellar and tarsal neurons. Although IR20a is not required for arginine-evoked calcium responses in tarsal GRNs, it is critical for arginine-driven feeding preference and the labellar PER. In contrast, IR20a is essential for low NaCl salt detection at both the cellular level and behavioral attraction. *UAS-hid* mediated genetic ablation of IR20a neurons further confirms the role of IR20a in arginine and low salt attraction (Fig. S1E - F). Thus, our data establish IR20a as a key component of both arginine and low NaCl salt attraction in *Drosophila*.

### IR20a functions with IR25a to detect arginine, while IR76b confers low NaCl salt responses

We next tested the requirement for IR25a and IR76b in *IR20a-GAL4* expressing tarsal GRNs for arginine and low-salt detection. Calcium imaging revealed that IR25a is essential for both arginine and low NaCl salt responses (Fig. 3A - C). In contrast, IR76b is dispensable for arginine detection but is required for NaCl responses (Fig. 3D - F). This suggests that IR20a forms a complex with IR25a to detect arginine, whereas low-salt sense requires the additional recruitment of the co-receptor IR76b.

**Figure 3.**
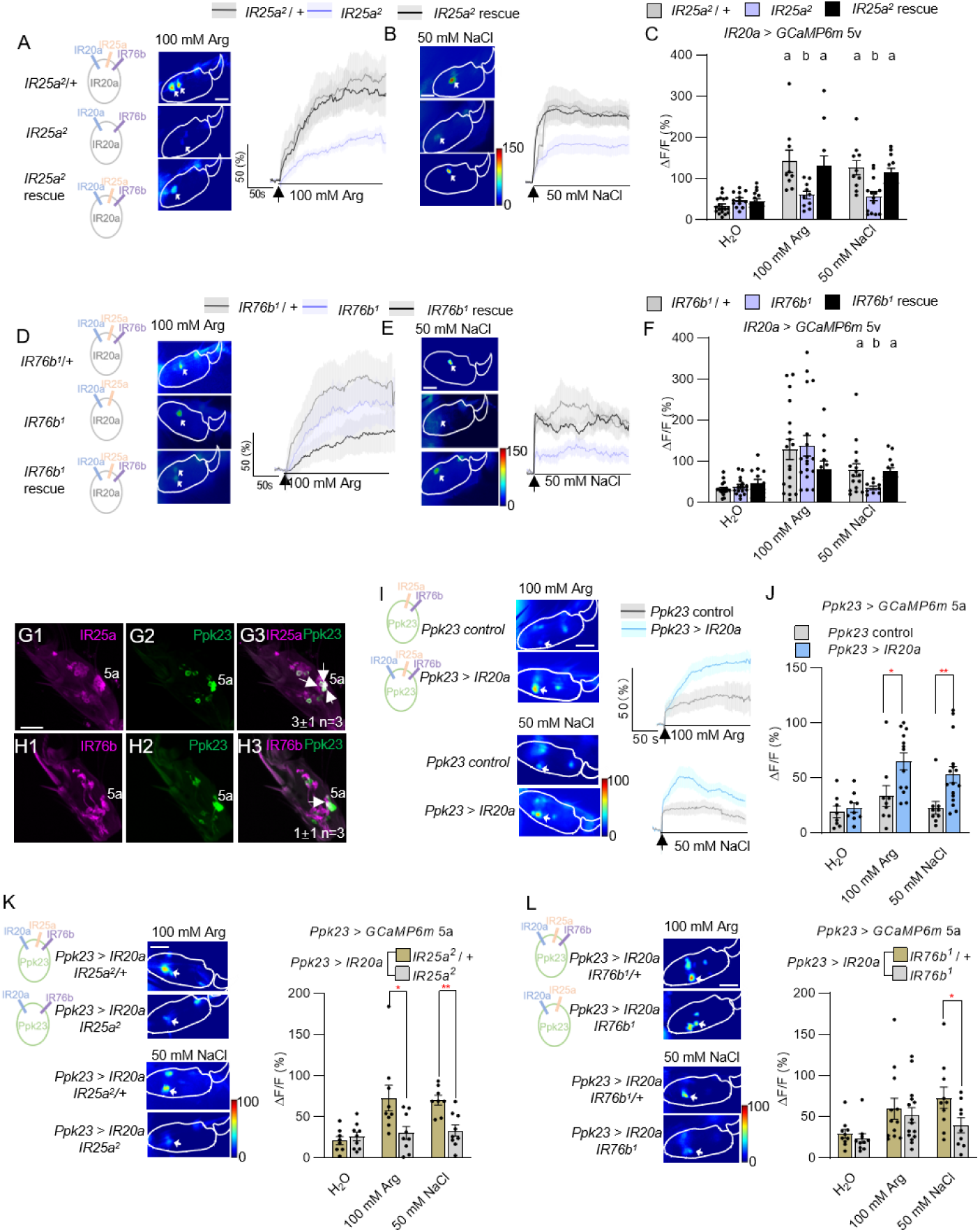
IR20a functions with IR25a to detect arginine, while IR76b is required for low-salt sensitivity. **(A - C)** IR25a is essential for both arginine and low-salt responses in IR20a tarsal GRNs. **(D - F)** IR76b is dispensable for arginine responses but is required for low-salt responses in IR20a tarsal GRNs. **(G)** Immunostaining for IR25a shows co-expression with *Ppk23-GAL4* in f5a sensilla; arrows indicate co-expression. **(H)** Co-localization of *Ppk23-GAL4* and *IR76b-QF* is restricted to a single cell body within the f5a sensilla; arrows indicate co-expression. **(I - J**) Ectopic expression of IR20a in *Ppk23-GAL4* tarsal 5a GRNs is sufficient to confer sensitivity to NaCl and arginine. **(K)** IR25a is necessary for the arginine and salt sensitivity conferred by IR20a ectopic expression in 5a GRNs. **(L)** IR76b is not required for the arginine sensitivity but is required for the salt sensitivity conferred by IR20a misexpression in 5a GRNs. Data represent mean ± SEM (n = 8 -18). In **(C)** and **(F)**, Kruskal-Wallis with Dunn’s post hoc test was used, different letters denote differences between designated groups (p < 0.05). In **(J - L)**, Mann-Whitney test was used, red asterisks indicate significant differences between indicated groups, *p < 0.05, **p < 0.01.

To determine if IR20a/IR25a is sufficient for arginine detection and IR20a/IR25a/IR76b for low-salt detection, we misexpressed IR20a in *Ppk23*-positive tarsal GRNs, which function in pheromone sensing and do not respond to ligands such as arginine or low NaCl salt (Fig. 3I) ^42^. Immunostaining confirmed that *Ppk23-GAL4* labels a pair of GRNs in tarsal 5a sensilla. All of these GRNs express IR25a, but only one pair also expresses IR76b (Fig. 3G - H). Calcium imaging of the *Ppk23* GRN in the 5a sensillum revealed that misexpression of IR20a conferred sensitivity to both arginine and NaCl (Fig. 3I - J). Crucially, mutation of *IR25a* abolished both the arginine and NaCl sensitivity acquired by IR20a misexpression (Fig. 3K). However, a mutation of *IR76b* only affected the conferred NaCl response but leaving arginine responses in tact (Fig. 3L). These data are consistent with our loss-of-function analysis and suggest that IR20a cooperates with IR25a to detect arginine, while the addition of IR76b is required to confer low-salt sensitivity.

To rigorously define the molecular mechanism, we characterized receptor ligand specificity by expressing different combinations of IR20a, IR76b, and IR25a in *Drosophila* S2 cells and performed calcium imaging. Expression of IR20a, IR25a, or IR76b alone, as well as co-expression of IR20a with IR76b or IR25a with IR76b, failed to elicit responses to arginine (Fig. 4A - F). In contrast, co-expression of IR20a with IR25a, or co-expression of all three genes elicited robust responses to 100 mM L-arginine (Fig. 4G - H). By contrast, neither IR20a nor IR25a alone, nor their co-expression, elicited responses to NaCl, whereas expression of IR76b alone introduced robust NaCl responses, and co-expression with either IR25a or IR20a abolished these responses (Fig. 4A, I -N). This is consistent with previous reports that IR76b forms a sodium channel in cultured cells ^4^, and further suggests that addition of either IR20a or IR25a into a IR76a-only complex suppresses NaCl channel activity. However, triple expression of IR20a, IR25a, and IR76b restored sensitivity to NaCl (Fig. 4A, O). To confirm the specificity of these responses, we tested all basic amino acids except for lysine at their maximum water-soluble concentrations using pooled stimulation paradigms. Neither the IR20a/IR25a complex nor the IR20a/IR25a/IR76b complex responded to any basic amino acid other than arginine (Fig. S2). Together, these results demonstrate that IR20a forms distinct receptor complexes with different co-receptor compositions, enabling multimodal detection of arginine and sodium.

**Figure 4.**
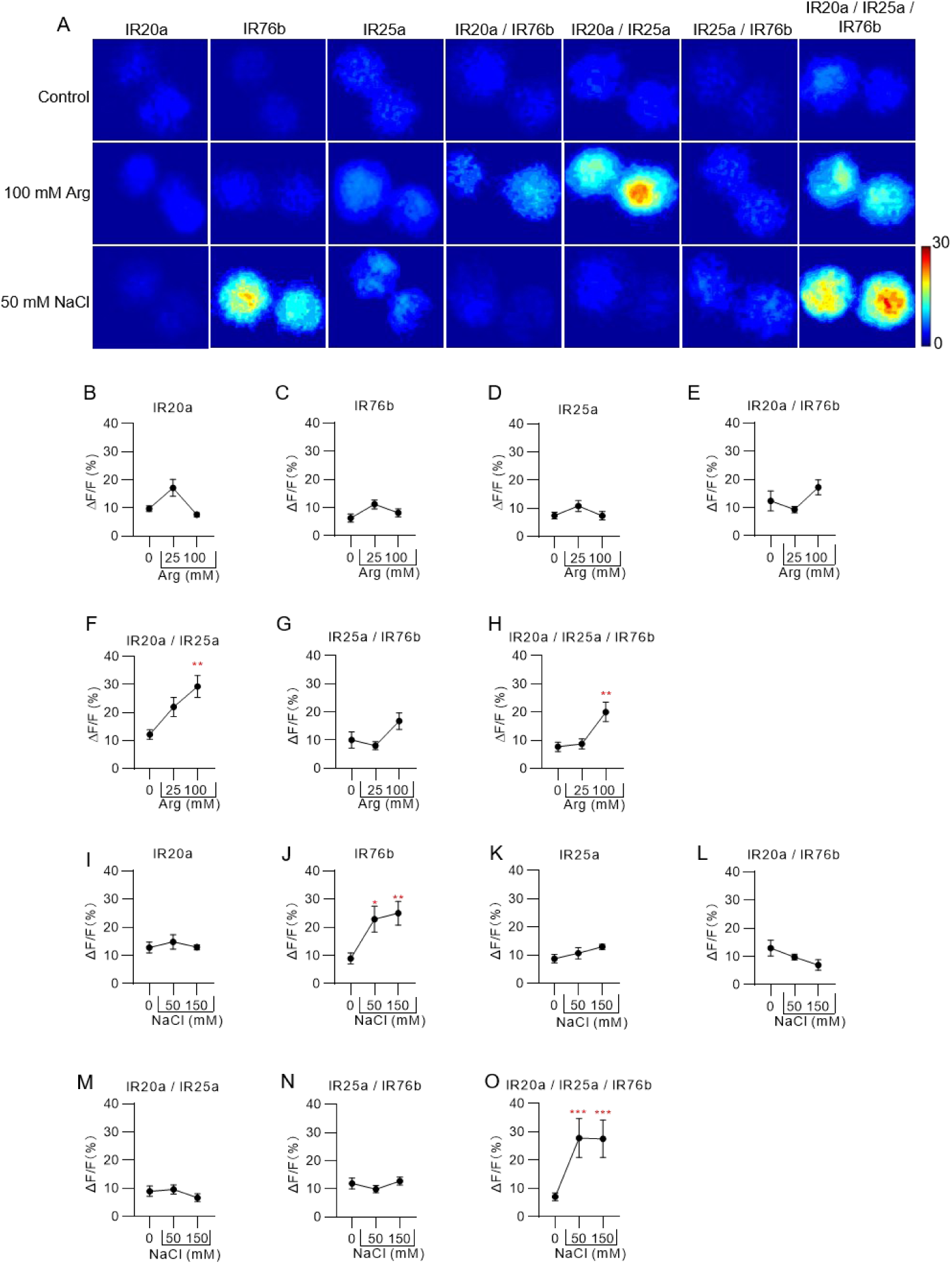
Co-expression of IR20a and IR25a confer arginine response, whereas adding IR76b provides low NaCl sensitivity in S2 cells. **(A)** Heat map of calcium responses in S2 cells transfected with various IR combinations. Only specific combinations confer sensitivity to arginine and NaCl. **(B - F)** Expression of IR20a, IR25a, or IR76b alone, as well as co-expression of IR20a with IR76b or IR25a with IR76b, failed to elicit arginine responses. **(G - H)** Co-expression of IR20a with IR25a, or triple express of IR20a, IR25a, and IR76b, confer arginine sensitivity. **(I - N)** Neither IR20a nor IR25a alone, nor their co-expression, elicited responses to NaCl, whereas expression of IR76b alone conferred low NaCl sensitivity. However, co-expression of IR76b with either IR20a or IR25a abolished NaCl sensitivity. **(O)** Triple expression of IR20a, IR25a, and IR76b confers NaCl response. Data represent mean ± SEM (n = 8 -17). Mann-Whitney test, *p < 0.05, **p < 0.01, ***p < 0.001.

### IR20a confers synergistic responses to an arginine and NaCl mixture

Given that amino acids and salt often occur together within a natural food source, the multimodal detection of arginine and NaCl salt by IR20a prompted us to test whether IR20a confers synergistic responses to their combination. PER assays with labellar stimulation revealed that a low concentration mixture (10 mM each) of arginine and NaCl elicits a stronger, IR20a-dependent response than either alone (Fig. 5A - B). To determine whether IR25a and IR76b are required co-receptors, we misexpressed IR20a in *Ppk23* tarsal 5a associated GRNs. Calcium imaging revealed that control neurons were unresponsive to a 10 mM mixture of arginine and NaCl (Fig. 5C), while IR20a misexpression conferred a robust response (Fig. 5C - D). This response was abolished in *IR25a* mutants but remained intact in *IR76b* mutants (Fig. 5E - F). Heterologous expression in S2 cells further confirmed the synergistic response. Specifically, cells co-expressing IR20a and IR25a did not respond to arginine or NaCl alone at concentrations of 10, 25, or 50 mM, but elicited a strong response to a 50 mM mixture (Fig. 5G - I and Fig. S3). However, the calcium response to the mixture was not further enhanced in the triple expression condition (Fig. 5 J - K). These results reveal that IR20a functions with IR25a to integrate arginine and low NaCl salt taste information to mediate synergistic responses, thereby enhancing feeding preference.

**Figure 5.**
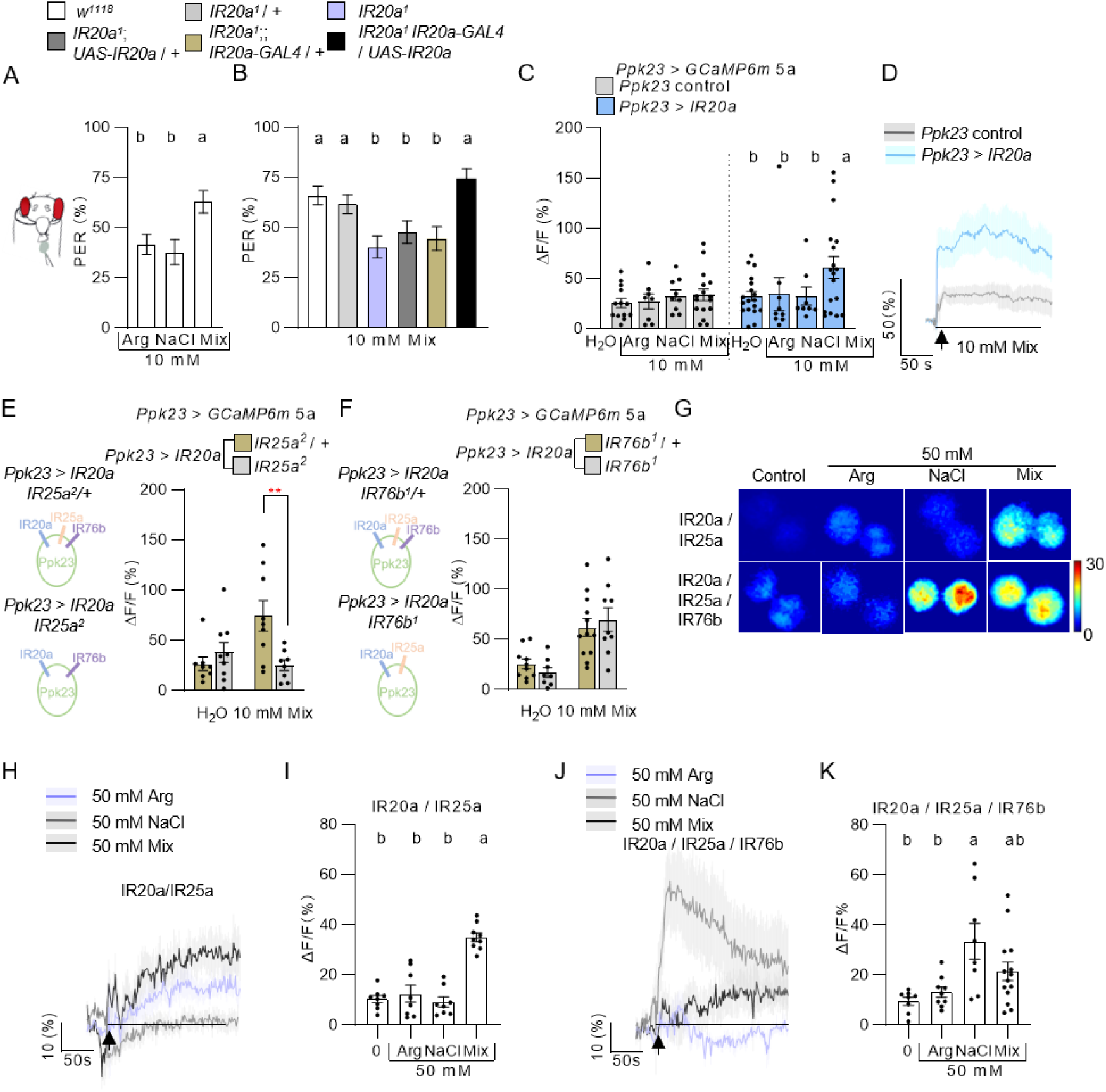
IR20a works with IR25a to mediate synergistic responses to arginine and NaCl. **(A - B)** PER assays reveal that a combination of arginine and NaCl elicits a synergistic response that is dependent on IR20a**. (C - D)** Calcium imaging shows that ectopic expression of IR20a in *Ppk23-GAL4* 5a GRNs confers the synergistic response to arginine-NaCl mixture. **(E - F)** Calcium imaging shows that IR25a, but not IR76b, is required for the synergistic response conferred by ectopic expression of IR20a in *Ppk23-GAL4* 5a GRNs. **(G - K)** Calcium imaging in S2 cells shows that co-expression of IR20a and IR25a confers a synergistic response to the arginine-NaCl mixture, whereas triple expression of IR20a, IR25a, and IR76b does not show a synergistic response to the arginine-NaCl mixture. Data represent mean ± SEM (n = 35 - 60 for PER assays, while n = 8 - 18 for calcium imaging). Kruskal-Wallis with Dunn’s post hoc test used in **(A - C, H, and J)**, Different letters denote differences between designated groups (p < 0.05), while Mann-Whitney test used in **(E - F)**, **p < 0.01.

### IR20a provides a tonic arginine and low NaCl salt signal parallel to the state-modulated IR56b pathway

Amino acid and salt preferences in flies are strongly influenced by internal nutritional state ^17,25,41,43^, but the sensory mechanisms underlying this modulation remain poorly understood. Having established the requirement of IR20a for arginine and low NaCl salt detection, we asked whether it mediates state-dependent shifts in preference. Two-choice feeding assays revealed that wild-type flies exhibit increased preference for arginine and low NaCl salt following amino acid deprivation and salt deprivation, respectively. However, these state-dependent changes occur independently of IR20a (Fig. 6A - B). Calcium imaging in *IR20a* tarsal GRNs confirmed that *IR20a* GRN responses to salt do not change between salt-fed and deprived conditions (Fig. 6C - D). In contrast, two-choice feeding assays showed that the state-dependent increase in salt preference is dependent on another low-salt tuning receptor, IR56b, which co-express with IR25a and IR76b in sweet GRNs but does not co-express with IR20a (Fig. 6E). Calcium imaging further confirmed IR56b is essential for state-dependent low salt sensitivity (Fig. 6F–G, and Fig. S4E). As a control, IR56b GRNs showed no change in sucrose and H2O sensitivity after salt deprivation (Fig. S4 A - D). These results suggest that IR20a provides a tonic arginine and low NaCl salt signal to maintain the stable detection of these essential nutrients, while the IR56b pathway adjusts sensitivity based on homeostatic need.

**Figure 6.**
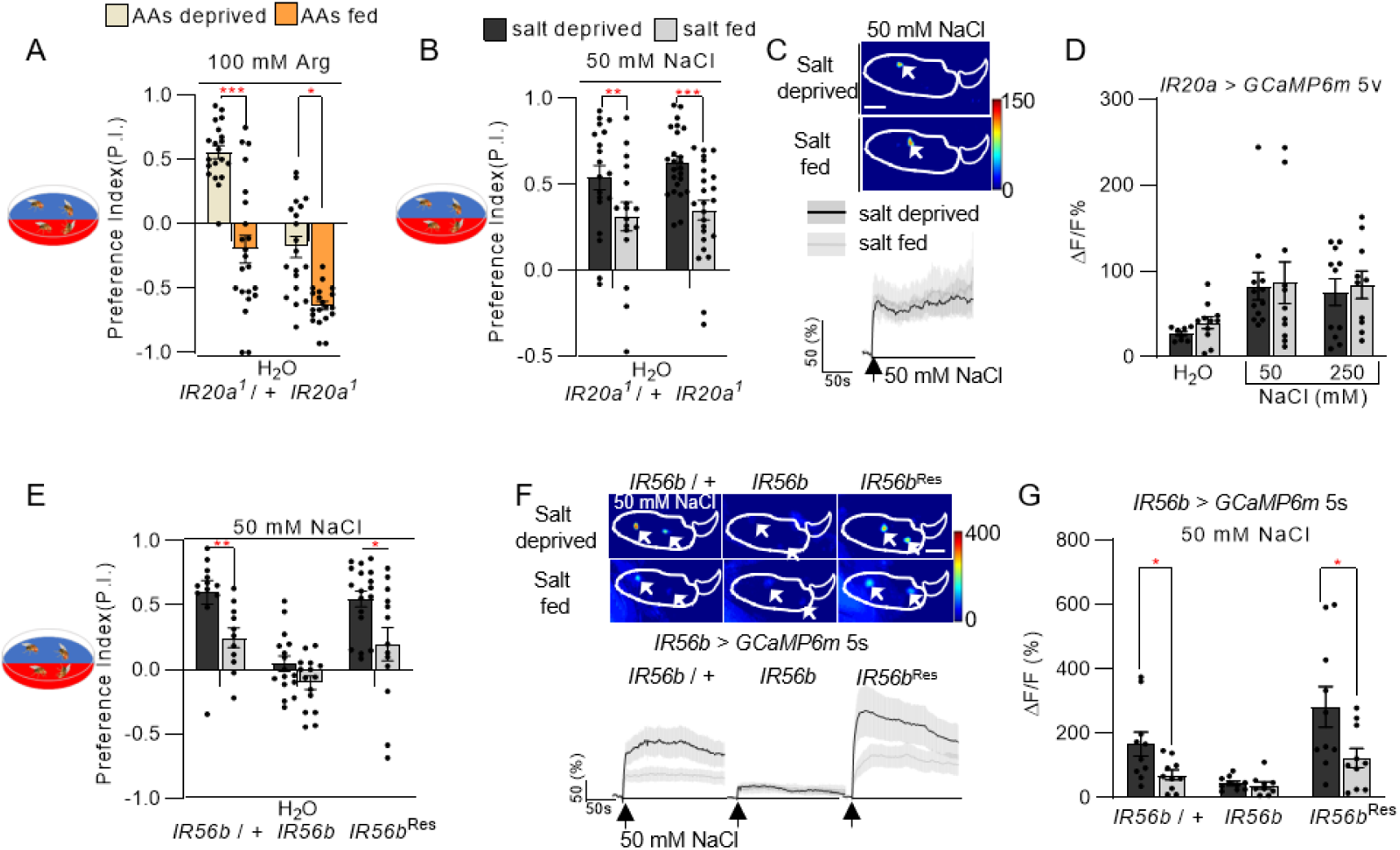
IR20a provides tonic sensitivity that cooperates with the state-modulated IR56b pathway. **(A)** Two-choice feeding assays show that amino acid deprivation increases preference for arginine, independent of IR20a. **(B)** Two-choice feeding assays show that salt deprivation increases preference for low NaCl, independent of IR20a. **(C - D)** Calcium imaging confirms that *IR20a-GAL4* tarsal GRN responses to NaCl are comparable in salt-deprived and salt-fed flies. **(E)** Two-choice feeding assays show that the state-dependent change in salt preference is dependent on IR56b. **(F - G)** Calcium imaging reveals that the state-dependent response of *IR56b* GRNs to NaCl requires IR56b. Data represent mean ± SEM (n = 11 - 31 for two-choice feeding assays, while n = 8 -12 for calcium imaging). Mann-Whitney test, *p < 0.05, **p < 0.01, ***p < 0.001.

## Discussion

In this study, we identify a specific population of taste sensory neurons in *Drosophila* that are broadly tuned to both low NaCl salt and specific amino acids arginine, implying that these essential nutrients are perceived through a multimeric channel containing IR20a, which, in combination with IR25a and IR76b, results in receptors with distinct preference for these ligands. To our knowledge, this represents the first evidence of signal integration between distinct taste modalities occurring at the level of the receptor complex itself.

### Refinement of Multimodal Coding in *Drosophila*

The complexity of the *Drosophila* gustatory system has long been underappreciated. While early models proposed a labeled-line organization similar to mammals, specifically humans and mice, comprehensive genetics, electrophysiological and calcium imaging studies have revealed that gustatory receptor neurons (GRNs) often exhibit broader tuning profiles than previously thought. For instance, sweet GRNs have been implicated in the detection of fatty acids and low salt, challenging the notion of exclusive modality coding ^17,20^.

Our identification of IR20a-expressing neurons significantly expands this perceptive range. Contrary to previous reports suggesting IR20a expression overlaps with amino-acid sensing sweet neurons ^37^, our intersectional genetic analysis reveals that IR20a marks a distinct population of non-sweet, non-bitter GRNs. While Ganguly et al suggested IR20a mediates detection of various amino acids (including glycine, serine, and phenylalanine) based on broad IR76b-GAL4 imaging and weak behavioral phenotypes ^37^. Our single-cell resolution imaging demonstrates that IR20a GRNs are narrowly tuned to arginine and proline, as well as monovalent salts (NaCl, KCl). Furthermore, while we find that tarsal responses to arginine are maintained in *IR20a* mutants, our behavioral assays establish that IR20a is strictly essential for the labellar-mediated preference for both arginine and low salt. This defines a novel, multimodal “savory” channel that is distinct from canonical single-modality sensory pathways.

### Combinatorial Assembly of IR Complexes

The IR gene family expanded during invertebrate evolution, presumably from a common ancestor of the iGluR gene family ^44,45^. The functional diversity of these receptors appears to rely on the combinatorial assembly of distinct subunits ^11,12,18,46^. Based on the work presented here, we propose that IR20a serves as a dual-function tuning receptor that dictates ligand specificity through differential complex formation.

Heterologous expression in S2 cells allows us to dissect this molecular logic. We observed that IR76b alone forms a constitutive Na^+^ channel, consistent with previous studies ^4^. However, co-expression with IR20a or IR25a abolishes this constitutive activity, suggesting that heteromeric assembly alters channel gating properties. We propose that the IR20a/IR25a complex forms the functional receptor for arginine, sufficient for ligand detection even in the absence of IR76b. In contrast, low NaCl salt detection requires the assembly of a tripartite complex: IR20a/IR25a/IR76b. *In vivo*, where these three subunits are co-expressed, it is likely that distinct stoichiometric populations, such as IR20a/IR25a dimers and IR20a/IR25a/IR76b multimers, coexist within the same dendritic membrane, thereby endowing the neuron with sensitivity to both amino acids and salts.

### Receptor-Level Signal Integration

Perhaps the most profound finding of this study is the mechanism of signal integration. Animals feeding in the wild encounter complex mixtures, and our behavioral data show that flies exhibit a synergistic preference for mixtures of arginine and NaCl. While synergy is typically attributed to central neural processing, our data indicate this integration occurs already peripherally at the gustatory receptor level, although additional modulation of this synergy in the brain is possible.

We demonstrate that the sensitivity to arginine of the IR20a/IR25a complex, which does not respond to NaCl alone, can be potentiated by addition of NaCl. This suggests that a possible allosteric mechanism where the binding of arginine to IR20a/IR25a exposes a latent binding site for Na^+^, or conversely, that Na^+^ stabilizes the ligand-bound conformation of the channel. This molecular integration offers a highly efficient mechanism for weighting and prioritizing nutrient signals; the preference for arginine over other amino acids is likely rooted in its unique physiological status, as it is one of the ten essential amino acids (EAAs) that *Drosophila* must acquire through diet, serving as the indispensable precursor for nitric oxide (NO) signaling and a limiting factor for oogenesis and fecundity ^47–49^. In the ecological niche of *Drosophila*, arginine and NaCl are concurrently enriched in yeast, their primary natural food source ^29^. By evolving a receptor complex that acts as a coincidence detector, the fly can specifically amplify the sensory output only when these two distinct nutrient classes are present together. This ensures that the IR20a pathway does not just signal the presence of any salt or any nitrogen source, but rather identifies high-quality, ecologically relevant resources that provide both essential amino acids and necessary electrolytes, thereby optimizing the fly’s foraging efficiency and reproductive success.

### Segregation of Tonic and State-Dependent Pathways

Finally, our results clarify the relationship between sensory detection and homeostatic regulation. Sodium salt preference in *Drosophila* is highly plastic, increasing during deprivation ^17^. We provide evidence that IR20a provides a tonic, stable signal for low NaCl salt and arginine that remains unchanged between salt-fed and salt-deprived conditions. In contrast, the state-dependent increase in NaCl salt preference is mediated by the previously identified IR56b pathway ^23,25^.

This parallel organization, one pathway (IR20a) ensuring the consistent recognition of vital nutrients and the other (IR56b) modulating sensitivity based on physiological need, represents a sophisticated survival strategy. It ensures that the fly never loses the ability to identify nutrient sources, even while dynamically adjusting its behavioral drive to consume them. In summary, IR20a and its co-receptors IR25a and IR76b represent a versatile molecular hub for multimodal nutrient detection and integration.

## Materials and Methods

### Fly husbandry and strains

Flies were reared on a standard cornmeal diet at 25°C and 70% humidity under a 12 hr light/12 hr dark cycle. The standard diet was prepared by dissolving the following ingredients in 10 L of water: 72.53 g agar, 520 g corn meal, 1100 g malt extract, 275 g yeast, 31.27 g propionic acid, and 14.07 g tegosept. A complete list of fly strains and source reagents used in this study is provided in the Key Resources Table.

### Molecular biology and transgenic flies

To generate *UAS-IR20a*, genomic DNA isolated from *w1118* flies served as the PCR template. The full genomic sequence of *IR20a* was amplified with primers listed in the Key Resources Table and inserted into the pUAST vector. All constructs were sequence-verified prior to embryo microinjection.

### Behavioral Analysis

#### Two-choice feeding assays

Assays were performed as previously described^19^ with minor modifications. Flies were 8–10 days old at the time of testing. For standard assays, newly eclosed flies were raised on standard food. For salt-manipulation experiments, newly eclosed flies were kept on standard food for 2–4 days and then transferred for 5 days to either salt-deficient food (0.7% agar, 563 mM sucrose) or salt-excess food (0.7% agar, 563 mM sucrose, 50 mM NaCl). 563 mM sucrose was selected to match the caloric density of standard food. For amino acid (AA) manipulation, flies were transferred after 2–4 days to either AA-deficient food (standard diet without yeast extract, supplemented with 16.5% maltose for isoenergetic balance) or AA-excess food (standard diet supplemented with 10% yeast extract and 4.16% maltose) for 5 days. Assays were conducted in Petri dishes (60 mm x 60 mm) containing 0.5% agarose mixed with the indicated tastants.

To visualize feeding, one half of the dish contained blue dye (brilliant blue FCF, 0.125 mg/ml) and the other contained red dye (sulforhodamine B, 0.2 mg/ml); dyes were swapped between conditions to control for color bias. Approximately 30 flies per vial were starved for 18–20 hours on wet filter paper, chilled on ice, and transferred to assay dishes in the dark for 2 hours. Feeding preference was scored by examining abdominal coloration under a stereomicroscope. The Preference Index (PI) was calculated as: PI = (N_color1_ – N_color2_) / N_total-fed_. Trials where >50% of flies failed to feed were excluded.

#### Proboscis Extension Reflex (PER)

PER assays were adapted from previous protocols ^19^. For leg stimulation, flies were anesthetized on ice and mounted dorsally on glass slides using double-sided tape, leaving appendages and the proboscis free. For labellar stimulation, flies were immobilized inside a white pipette tip with only the head and labellum exposed. 20–30 flies were mounted per slide and allowed to recover for 1 hour at 25°C (70% humidity). Prior to testing, flies were water-satiated until they ceased drinking. Stimulation was performed by touching the legs or labellum with the indicated tastant (NaCl or amino acids) for 3–5 seconds. A positive response was recorded if full proboscis extension occurred. Each fly was also tested with 20 mM sucrose as a positive control. Two trials were conducted per tastant with a > 20-minute inter-trial interval. Responses were scored as 100% (extension in both trials), 50% (extension in one trial), or 0% (no extension). Flies failing to respond to the sucrose control in both trials were excluded from the analysis.

### Immunohistochemistry

Dissection and staining were performed on 6–10 day-old flies. Flies were CO2-anesthetized, and labella and legs were dissected using a razor blade; legs were severed between the third and fourth tarsal segments. Tissues were immediately immersed in 4% paraformaldehyde and briefly centrifuged (1000 rpm, 30 s) to ensure complete submersion. Following fixation on ice for 1 hour, tissues were washed three times (15 min each) in PBST (1x PBS + 0.3% Triton X-100) at room temperature. Blocking was performed in PBST containing 5% horse serum for 2 hours at room temperature. Tissues were incubated with primary antibodies in blocking solution overnight at 4°C, followed by three 15-minute washes in PBST. Tissues were then re-blocked in PBST + 5% horse serum for 30 minutes and incubated with secondary antibodies for 3 hours at room temperature. After three final washes in PBST, samples were mounted in antifade medium and sealed with nail polish. Images were acquired on a Zeiss LSM 800 confocal microscope using 20X or 40X objectives.

#### Primary antibodies

Chicken anti-GFP (1:2000), rabbit anti-RFP (1:2000), mouse anti-HA (1:500), mouse anti-nc82 (1:5000), and rabbit anti-IR25a (1:200). Anti-IR25a was generated against synthetic peptides (SKAALRPRFNQYPATFKPRF and DVAEANAERSNAADHPGKLVDGV) as described in Benton et al ^44^.

#### Secondary antibodies

Goat anti-chicken Alexa 488 (1:2000), goat anti-rabbit Alexa 488 (1:200), goat anti-mouse Alexa 568 (1:2000), goat anti-rabbit Alexa 568 (1:2000), and goat anti-mouse Alexa 647 (1:2000).

### Calcium Imaging

#### Single GRN calcium imaging

single tarsal GRN calcium imaging was performed as previously described ^40^. *UAS-GCaMP6m* was driven by specific GAL4 lines in 7–10 day-old females. Legs were prepared by severing at the tibia-femur junction. The cut end was sealed with grease and attached to double-sided tape on a glass-bottom dish (35 x 35 mm), leaving the 4th and 5th tarsal segments suspended. Warm agarose (50–60°C) was applied to the tape to stabilize the preparation. Imaging was performed on a Nikon Ni-U upright microscope with a 40X water immersion objective. Fluorescence changes were monitored in cell bodies relative to adjacent background regions. Recording began 10 seconds prior to tastant application and continued for 2–3 minutes. The response was quantified as DF/F (%) = (F - F_0_) / F_0_, where F_0_ is the mean baseline fluorescence of the 5 seconds preceding stimulation. The maximum DF/F value was used for analysis.

### S2 cell culture and imaging

S2 cell culture and imaging were performed by adapted protocols from previous ^24^. *Drosophila* S2 cells were cultured in Sf-900 II SFM supplemented with 10% FBS and 0.5% penicillin/streptomycin at 25°C. Cells were transfected with pAc5.1 vectors encoding IR20a (genomic), IR25a (cDNA), IR76b (cDNA), and GCaMP6m. Imaging was conducted using a Nikon Ni-U microscope (10× objective). Prior to imaging, culture medium was replaced with isotonic bath buffer (10 mM HEPES, 10 mM glucose, 250 mM NMDG-Cl; pH 7.4; 520 mOsm). After a 2-minute equilibration, the buffer was removed to a minimal volume. Following a 10-second baseline recording (F_0_), cells were stimulated with ligand solutions containing the test tastant (NaCl or arginine). To maintain constant osmolarity (520 mOsm) across all conditions, the concentration of NMDG-Cl in the stimulation buffer was adjusted inversely to the tastant concentration. Fluorescence was recorded for 120 seconds, and background-subtracted peak DF/F (%) responses were quantified.

### Quantification and Statistical Analysis

Normality of data distribution was assessed prior to test selection. For non-normally distributed data, comparisons between two groups were performed using the Mann-Whitney test, while multiple group comparisons were analyzed using the Kruskal-Wallis test followed by Dunn’s post-hoc test. Data are presented as mean standard error of the mean (SEM). The value n represents the number of preparations for immunostaining, the number of neurons for calcium imaging, the number of flies for PER, or the number of plates for two-choice feeding assays. Specific statistical details are provided in Dataset S1 - S8.

## KEY RESOURCES TABLE

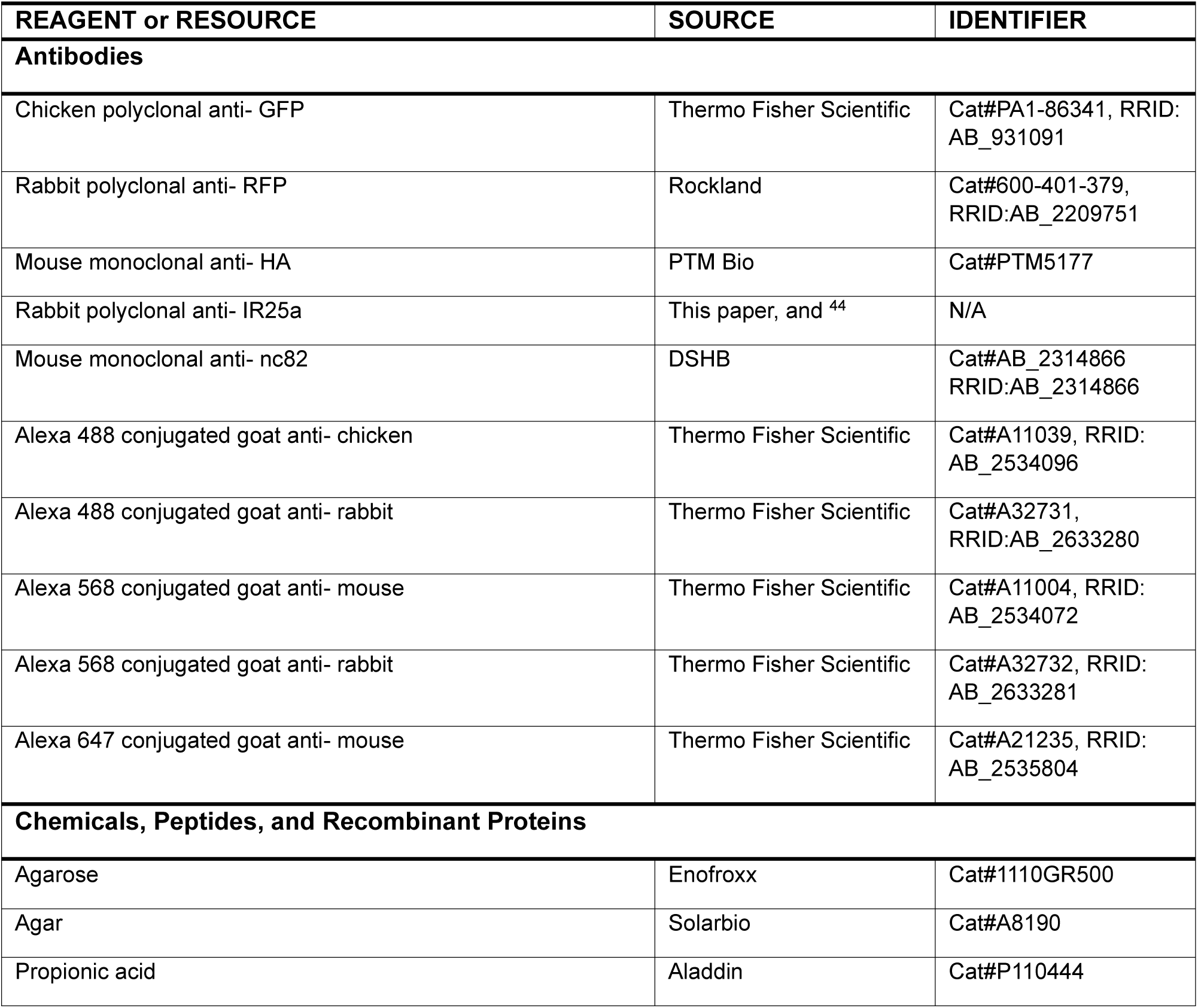
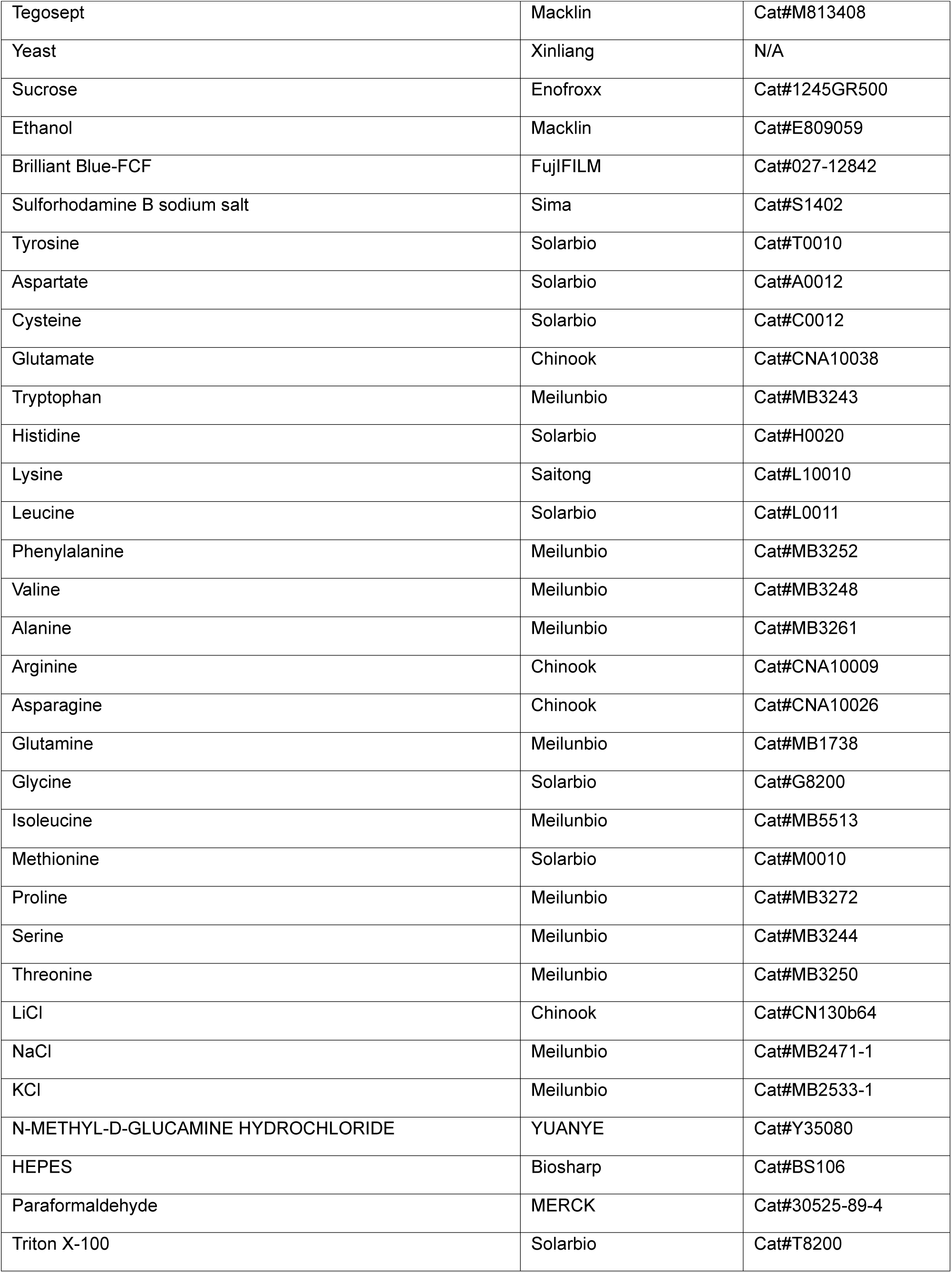
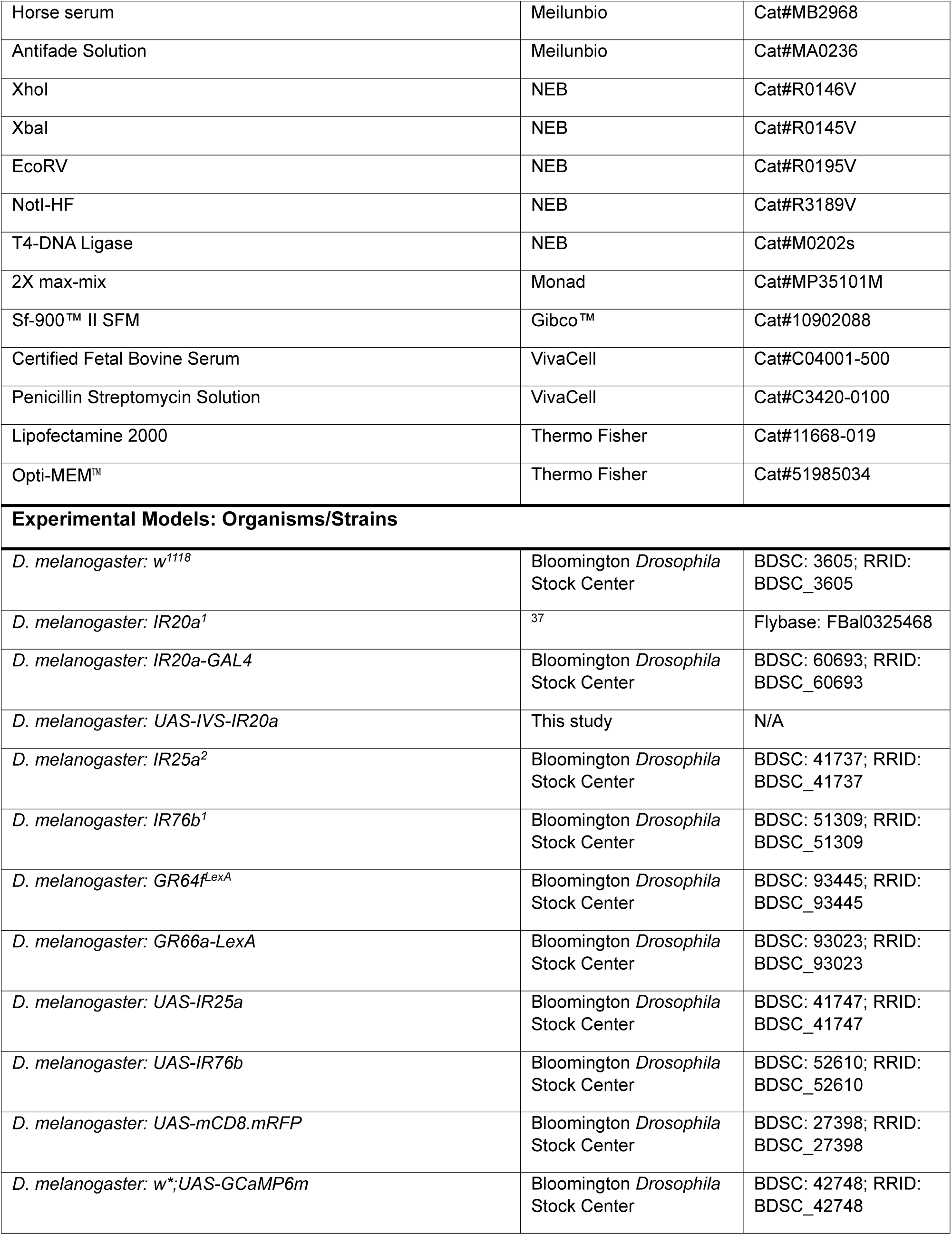
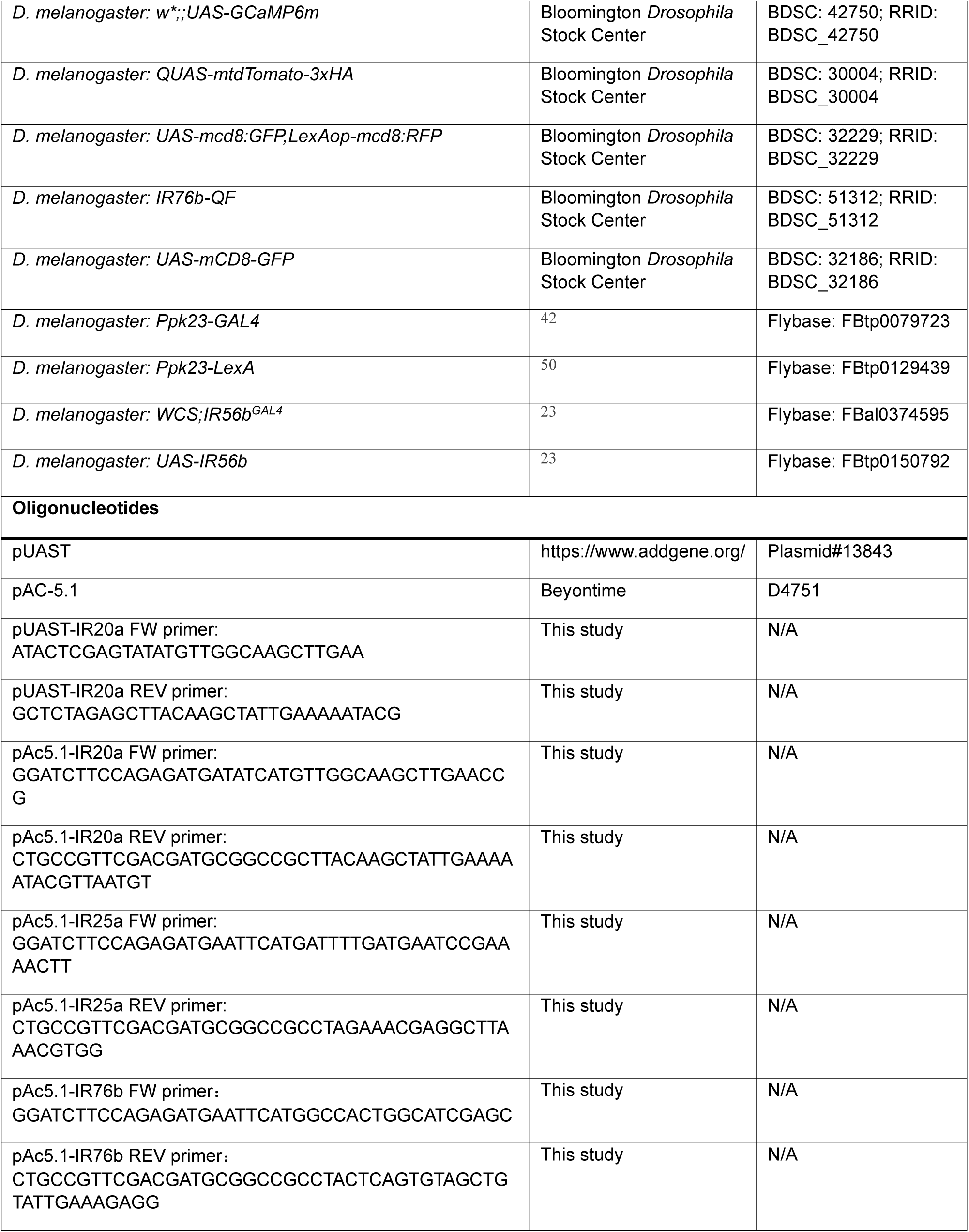
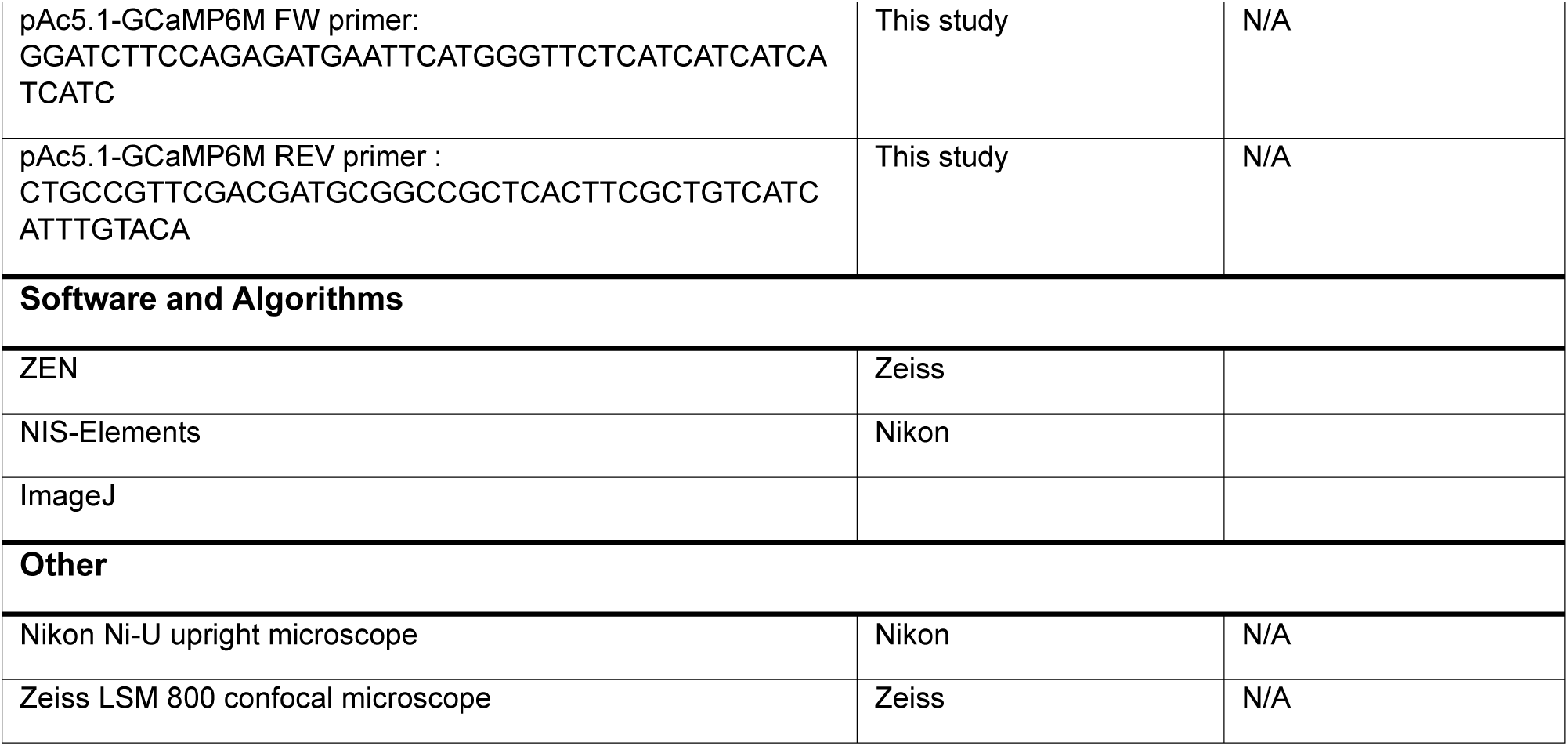

## Supporting information

Data S1-Genotypes and statistical details for Fig.1

Data S2-Genotypes and statistical details for Fig.2

Data S3-Genotypes and statistical details for Fig.3

Data S4-Genotypes and statistical details for Fig.4

Data S5-Genotypes and statistical details for Fig.5

Data S6-Genotypes and statistical details for Fig.6

Data S7-Genotypes and statistical details for Fig.S1

Data S8-Genotypes and statistical details for Fig.S2

Data S9-Genotypes and statistical details for Fig.S3

Data S10-Genotypes and statistical details for Fig.S4

## Acknowledgments

We thank Dr. Minjiao Liu for making the *pUAST-IR20a* construct. We thank Drs. Hui Xiao, Guoqiang Zhang, the Bloomington *Drosophila* Stock Center, the Tsinghua Fly Center, and VDRC for fly strains and plasmids, and the Animal Center of Yunnan University for antibodies. We thank Prakash Pandey for his critical comments on the manuscript. This work was supported by grants from the National Natural Science Foundation of China (grant no. 32070997 to Y.C), Applied Basic Research Foundation of Yunnan Province (grant no. 2019FY003018 to Y.C.), Research Startup Funding of Yunnan University (to Y.C.) and the NIH (1RO1GMDC05606-01 to H.A.)

## Authors contribution

B.W. performed the immunostaining, leg calcium imaging, S2 cell culture and imaging, Proboscis extension reflex assays and two-choice feeding assays, G.Q. performed immunostaining, leg calcium imaging, and two-choice feeding assays, X.W. performed Proboscis extension reflex assays. B.W., G.Q., and Y.C. analyzed data, H.A. and Y.C. conceived and supervised the project, designed experiments, and wrote the paper.

---

**Figure S1.**
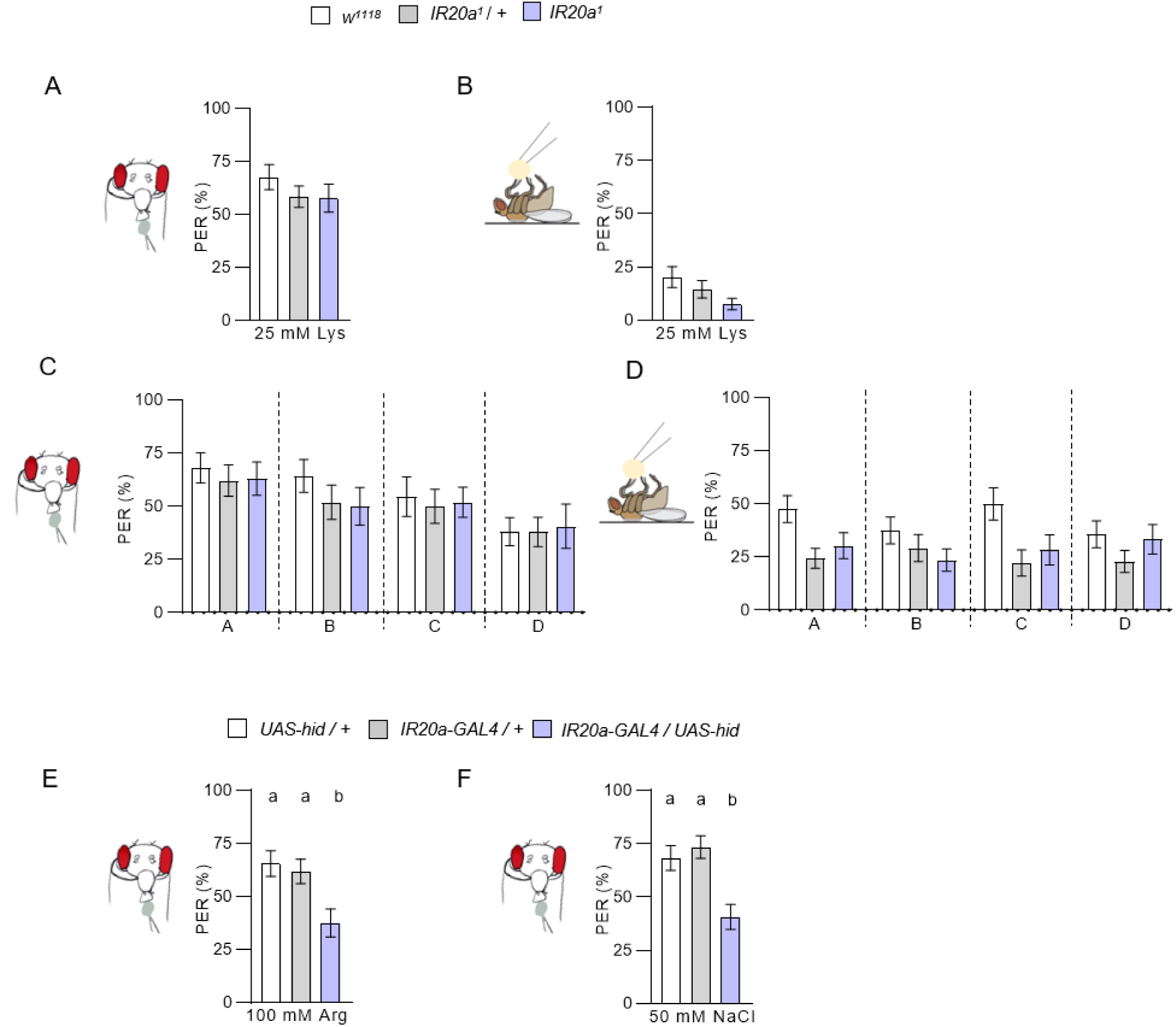
*IR20a* is specifically required for arginine and NaCl attraction. **(A - B)** *IR20a* is not required for 25 mM lysine preference in PER assays. (n = 18 – 35). **(C - D)** *IR20a* is not required for preference in amino acid mixture in PER assays, n = 16 - 40. **Group A**: 25 mM each of Ser, Thr, and Phe. **Group B**:100 mM Met, 75 mM Ile, 50 mM Val and Leu, 25 mM Trp. **Group C**:100 mM Gly, 75 mM Ala, 50 mM His, 25 mM Cys. **Group D**: 25 mM Glu, 50 mM Gln, 50 mM Asn, 25 mM Asp, 0.5 mM Tyr. **(E)** Ablation of *IR20a-GAL4* by *UAS-hid* reduces PER response to both arginine and NaCl. Data represent mean ± SEM (n = 38 - 49). Kruskal-Wallis with Dunn’s post hoc test, different letters denote differences between designated groups (p < 0.05).

**Figure S2.**
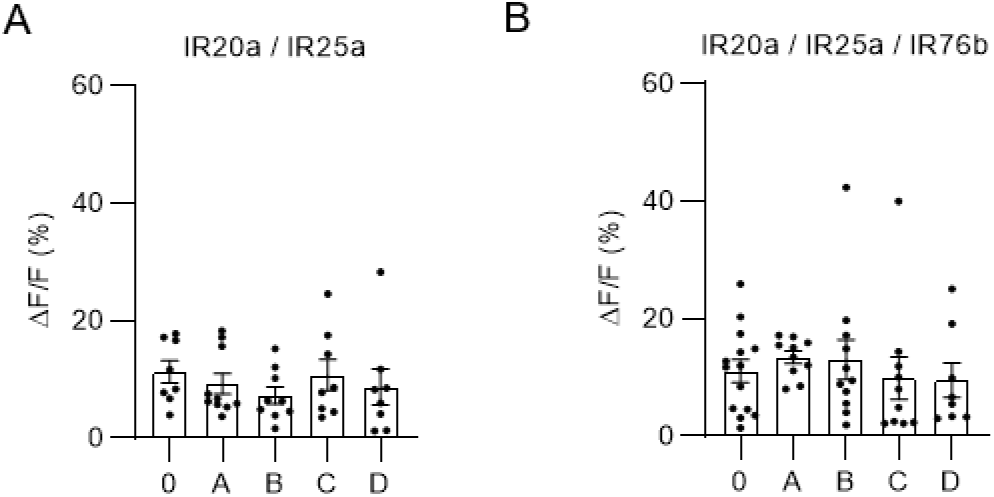
Neither co-expression of IR20a and IR25a nor triple expression of IR20a, IR25a, and IR76b in S2 cells confers broad amino acid sensitivity. **(A - B)** Calcium imaging shows that heterologous expression of IR20a and IR25a, or triple expression of IR20a, IR25a, and IR76b in S2 cells does not elicit responses to various amino acid mixtures. **Group A**: 25 mM each of Ser, Thr, and Phe. **Group B**:100 mM Met, 75 mM Ile, 50 mM Val and Leu, 25 mM Trp. **Group C**:100 mM Gly, 75 mM Ala, 50 mM His, 25 mM Cys. **Group D**: 25 mM Glu, 50 mM Gln, 50 mM Asn, 25 mM Asp、0.5 mM Tyr. Data represent mean ± SEM (n = 8 -14).

**Figure S3.**
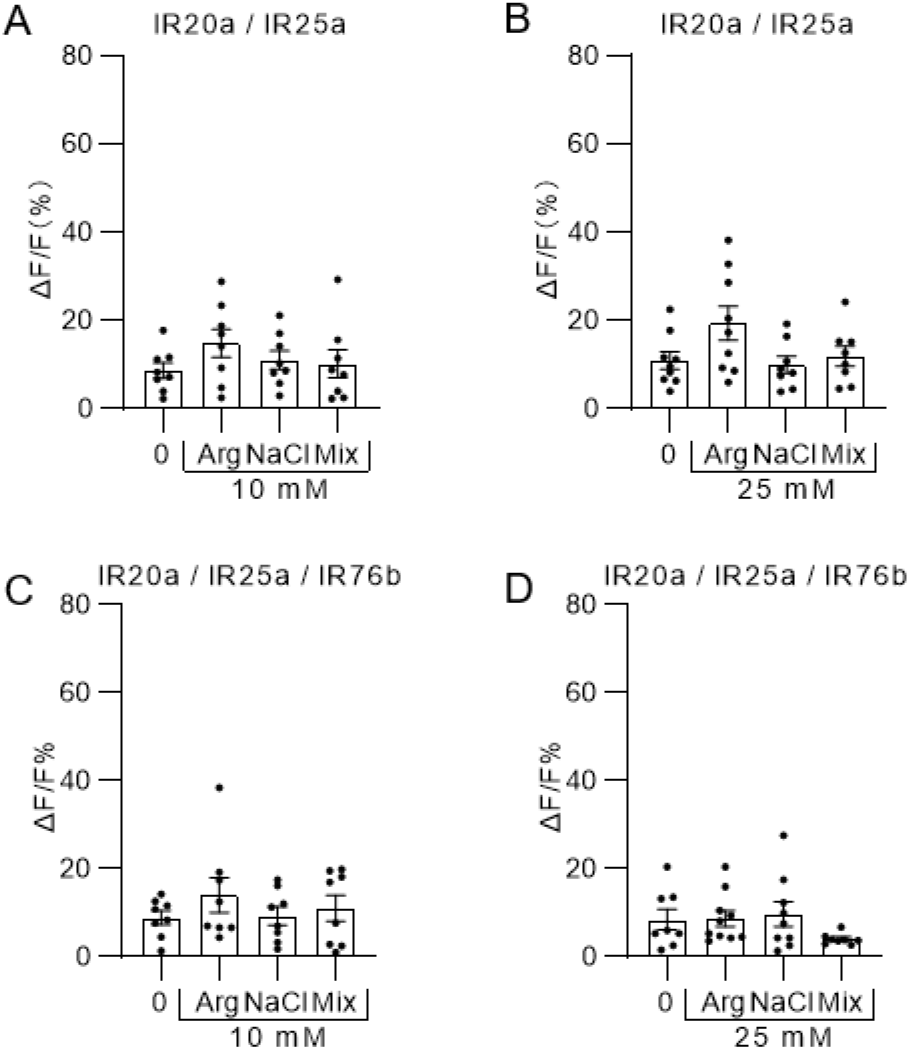
Synergistic responses to an arginine and NaCl mixture in S2 cells. **(A–D)** Calcium imaging shows that heterologous expression of IR20a and IR25a, or triple expression of IR20a, IR25a, and IR76b in S2 cells does not elicit synergistic responses to 10 or 25 mM of arginine-NaCl mixture. Data represent mean ± SEM (n = 8 -10).

**Figure S4.**
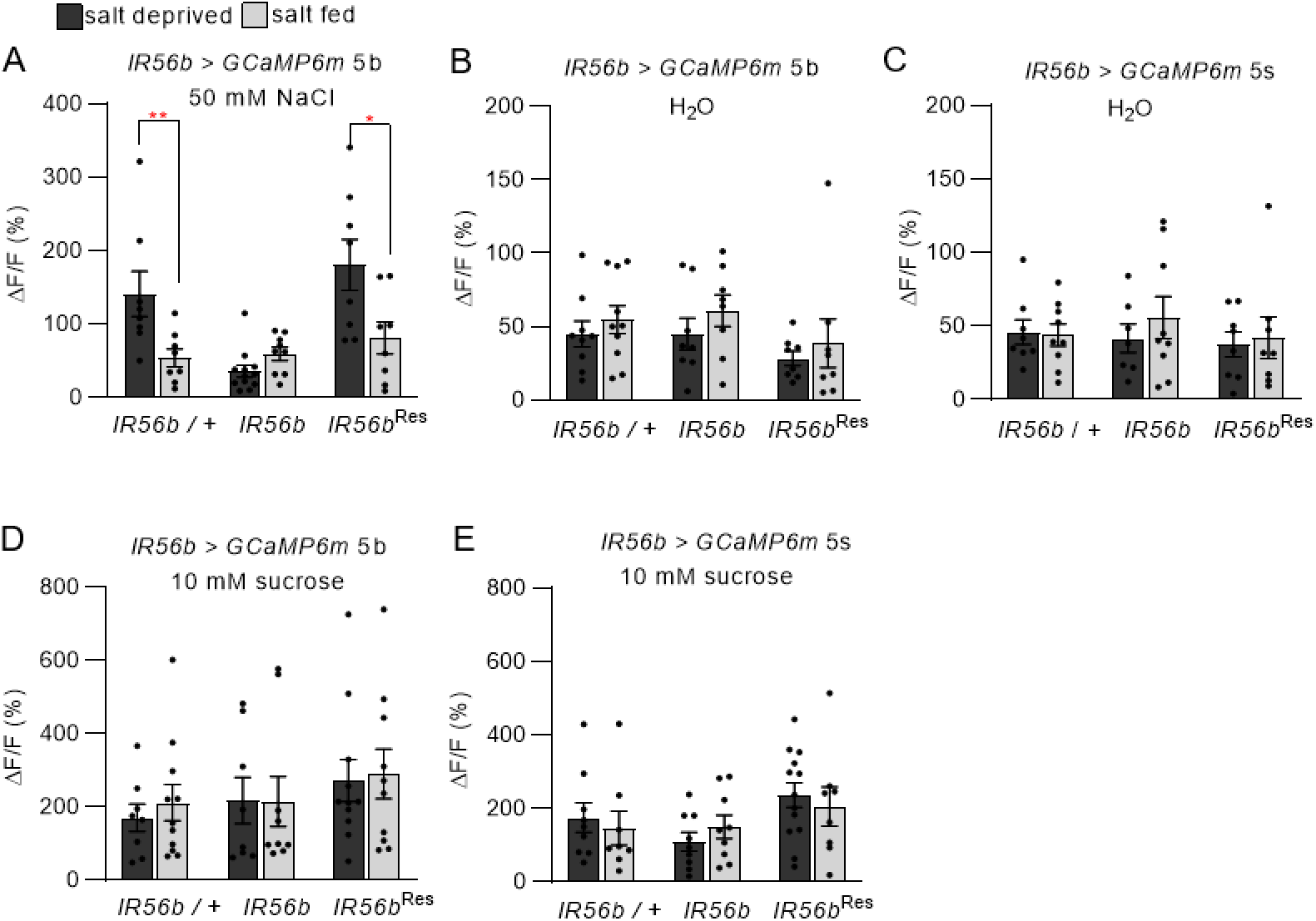
IR56b is required for state-dependent low NaCl response. **(A)** The state-dependent increase in salt preference is abolished in *IR56b* mutants, and this phenotype is rescued by reintroducing IR56b. **(B - C)** Calcium respond to H_2_O in *IR56b* controls, mutants and rescue. **(D - E)** Calcium responses to sucrose are independent on salt-fed condition in wild-type control, *IR56b* mutants and rescue. Data represent mean ± SEM (n = 8 - 13). Mann-Whitney test, *p < 0.05.

